# Evolution of Nodal and Nodal-related genes and the putative composition of the heterodimer that triggers the Nodal pathway in vertebrates

**DOI:** 10.1101/483974

**Authors:** Juan C. Opazo, Shigehiro Kuraku, Kattina Zavala, Jessica Toloza-Villalobos, Federico G. Hoffmann

**Author notes:** Corresponding author: Juan C. Opazo, Instituto de Ciencias Ambientales y Evolutivas, Facultad de Ciencias, Universidad Austral de Chile, Valdivia, Chile.

## Abstract

Nodal is a signaling molecule that belongs to the transforming growth factor-beta superfamily that plays key roles during the early stages of development of animals. Nodal forms an heterodimer with a GDF1/3 protein to activate the Nodal pathway. Vertebrates have a paralog of nodal in their genomes labeled Nodal related, but the evolutionary history of these genes is a matter of debate, mainly because of variable numbers of genes in the vertebrate genomes sequenced so far. Thus, the goal of this study was to investigate the evolutionary history of the Nodal and Nodal-related genes with an emphasis in tracking changes in the number of genes among vertebrates. Our results show the presence of two gene lineages (Nodal and Nodal-related) that can be traced back to the ancestor of jawed vertebrates. These lineages have undergone processes of differential retention and lineage-specific expansions. Our results imply that Nodal and Nodal-related duplicated at the latest in the ancestor of gnathostomes, and they still retain a significant level of functional redundancy. By comparing the evolution of the Nodal/Nodal-related with GDF1/3 gene family, it is possible to infer that there are at least four types of heterodimers that can trigger the Nodal pathway among vertebrates.

## 1 INTRODUCTION

Understanding the evolution of genes that are involved in early developmental processes represents an important challenge in biology (Gilbert, 2017). In most of the cases, the information that is available is restricted to the presence of a particular developmental pathway, and of the genes that participate, but much less is known regarding the evolutionary history of the genes and pathways (Jones et al., 1995; Levin et al., 1995; Seleiro et al., 1996; Joseph and Melton, 1997; Raya et al., 2004a; Lagadec et al., 2015; Tadjuidje et al., 2016; Li et al., 2017).

The gene Nodal encodes for a signaling molecule that belongs to the transforming growth factor-beta superfamily (TGF-β). It is a single-copy gene in most animal groups, but in vertebrates it has a paralog called Nodal-related. The Nodal pathway is active during the pre-gastrulation and gastrulation stages of development (Shen, 2007; Zinski et al., 2018) and is responsible for multiple fundamental processes such as mesoderm induction and patterning, endoderm function, neural patterning, establishment of left-right asymmetry, dorsal-ventral axis specification, anterior-posterior axis formation and maintenance of undifferentiated embryonic stem cells (Conlon et al., 1994; Jones et al., 1995; Feldman et al., 1998; Schier, 2003; Shen, 2007). Recently, three independent groups have shown that Nodal needs to form heterodimers with the GDF1/3 protein, encoded by another member of the TGF-β superfamily, to activate the signaling pathway (Bisgrove et al., 2017; Montague and Schier, 2017; PellicciaJ.L. et al., 2017). To gain insights into the evolutionary history of the Nodal pathway, Opazo & Zavala (2018) studied the evolutionary history of the GDF1/3 gene family and found that the vast majority of vertebrates have a single GDF1/3 ortholog in their genome. The two exceptions are amphibians and mammals, in which lineage-specific duplications gave rise the separate GDF1 and 3 genes: GDF1_A_ and 3_A_ for amphibians and GDF1_M_ and 3_M_ for mammals.

On the other hand, the evolution of its partner, the Nodal gene and its paralog, nodal-related, remains poorly understood. There is extensive variation in the number of genes in the Nodal gene family among vertebrates, ranging from one in mammals and some sauropsids to six in amphibians (e.g., *Xenopus laevis*). In fact, most vertebrate genomes include copies of Nodal and Nodal-related, but their phyletic distribution is not clear, and nomenclature is problematic, as different names have been applied to orthologs, and conversely, the same name has been applied to paralogous copies. In addition, the evolutionary history of the Nodal gene and its homologs is a matter of debate mainly because the number of Nodal and Nodal-related genes lineages present in vertebrates is not well defined, and sampling differences complicate comparisons among studies (Fan and Dougan, 2007; Kuraku and Kuratani, 2011).

Accordingly, the goal of this study was to reconstruct the evolutionary history of the Nodal and Nodal-related genes in vertebrates to 1) define the number of Nodal and Nodal-related genes in vertebrates, 2) reconstruct evolutionary relationships among the vertebrate Nodal and Nodal-related to unravel the duplicative history of the genes, with special attention to amphibians and teleost fish, and 3) integrate the history of the Nodal and GDF1/3 gene families to gain insights into the evolution of the heterodimer that is responsible for triggering the Nodal pathway. Our results indicate that the duplication giving rise to Nodal and Nodal-related traces back at least to the last common ancestor of jawed vertebrates. These two genes have undergone differential retention and lineage-specific expansions, where the lack of Nodal orthologs in birds, crocodiles, turtles and snakes, as well as, the expansion of Nodal-related genes in anurans represent the most salient results. Finally, by comparing the presence and absence of the Nodal/Nodal-related and GDF1/3 gene family members in different lineages, we infer that at least four different types of heterodimers can activate the Nodal pathway among vertebrates.

## 2 MATERIALS AND METHODS

### 2.1 Sequence data

We implemented bioinformatic searches to retrieve Nodal and Nodal-related genes in mammals, birds, reptiles, amphibians, coelacanths, teleost fish, holostean fish, cartilaginous fish, cyclostomes, urochordates and cephalochordates (Supplementary Table S1). We identified pieces containing Nodal and Nodal-related genes in the Ensembl database (Zerbino et al., 2018) using BLASTN or National Center for Biotechnology Information (NCBI) database (refseq_genomes, htgs, and wgs) (Geer et al., 2010) using TBLASTN (Altschul et al., 1990) with default settings. Conserved synteny was also used to define the genomic region containing Nodal and Nodal-related genes. Once identified, genomic pieces were extracted including the 5′and 3′ flanking genes. After extraction, we curated the existing annotation by comparing known exon sequences to genomic pieces using the program Blast2seq with default parameters (Tatusova and Madden, 1999). Putatively functional genes were characterized by an open intact reading frame with the canonical exon/intron structure typical of Nodal/Nodal-related gene. Sequences derived from shorter records based on genomic DNA or cDNA were also included to attain a broad taxonomic coverage.

### 2.2 Phylogenetic analyses

Amino acid sequences were aligned using the L-INS-i strategy from MAFFT v.7 (Katoh and Standley, 2013). We used the proposed model tool of IQ-Tree v1.6 (Trifinopoulos et al., 2016) to select the best-fitting model of amino acid substitution (JTT + F + R6). We employed the maximum-likelihood method to obtain the best tree using the program IQ-Tree v1.6 (Trifinopoulos et al., 2016); support for the nodes was assessed with 1,000 bootstrap pseudoreplicates using the ultrafast routine. Bayesian searches were conducted in MrBayes v.3.1.2 (Ronquist and Huelsenbeck, 2003). Two independent runs of six simultaneous chains for 1 × 10^7^ generations were set, and every 1,000 generations were sampled using default priors. The run was considered to have reached convergence once the likelihood scores formed an asymptote and the average standard deviation of the split frequencies remained < 0.01. We discarded all trees that were sampled before convergence, and we evaluated support for the nodes and parameter estimates from a majority rule consensus of the last 5,000 trees. Human and opossum growth differentiation factor 10 (GDF10) and bone morphogenetic protein 2 (BMP2) sequences were used as outgroups. Divergence times were obtained from the TimeTree server (Kumar et al., 2017).

### 2.3 Assessment of Conserved Synteny

We examined genes found upstream and downstream of the Nodal and Nodal-related genes of the species below mentioned to encompass diverse vertebrate lineages. For comparative purposes, we used the estimates of orthology and paralogy derived from the EnsemblCompara database (Herrero et al., 2016); these estimates are obtained from an automated pipeline that considers both synteny and phylogeny to generate orthology mappings. These predictions were visualized using the program Genomicus v94.01 (Louis et al., 2015). Our assessments were performed in humans (*Homo sapiens*), chicken (*Gallus gallus*), American alligator (*Alligator mississippiensis*), Chinese softshell turtle (*Pelodiscus sinensis*), anole lizard (*Anolis carolinensis*), Western clawed frog (*Xenopus tropicalis*), axolotl (*Ambystoma mexicanum*), coelacanth (*Latimeria chalumnae*), spotted gar (*Lepisosteus oculatus*), zebrafish (*Danio rerio*), and *Callorhinchus milii*, the elephant fish, which is sometimes labeled as elephant shark, and inshore hagfish (*Eptatretus burgeri*). In the case of the elephant fish, American alligator and axolotl, flanking genes were examined using the Entrez gene database from the NCBI (Maglott et al., 2005).

## 3 RESULTS

### 3.1 Identification of two Nodal paralogs in vertebrates

We recovered the monophyly of the Nodal and Nodal-related sequences from chordates with strong support in our Bayesian and ML analyses (Fig. 1). These analyses place the single ortholog of lancelets (cephalochordates) and sea squirts (urochordates) as sister to the clade that includes the Nodal and Nodal-related sequences of vertebrates (Fig. 1). The Nodal genes of jawed vertebrates were placed in two strongly supported clades, corresponding to the Nodal and Nodal-related genes respectively (Fig. 1). The Nodal clade includes cartilaginous fish, holostean and teleost fish, coelacanth, amphibians, lizards and mammals, and the Nodal-related clade includes cartilaginous fish, holost and teleost fish, coelacanth, amphibians, and sauropsids (Fig. 1). The Nodal-like genes of cyclostomes are placed in a strongly supported monophyletic group sister to the Nodal gene of jawed vertebrates, although support for this node is weak (Fig. 1). The Nodal gene is absent from the genomes of snakes, turtles, crocodilians, and birds, whereas the Nodal-related gene is absent from the genomes of all mammals examined. In all cases, deviations from the expected organismal phylogeny were weakly supported.

**Figure 1.**
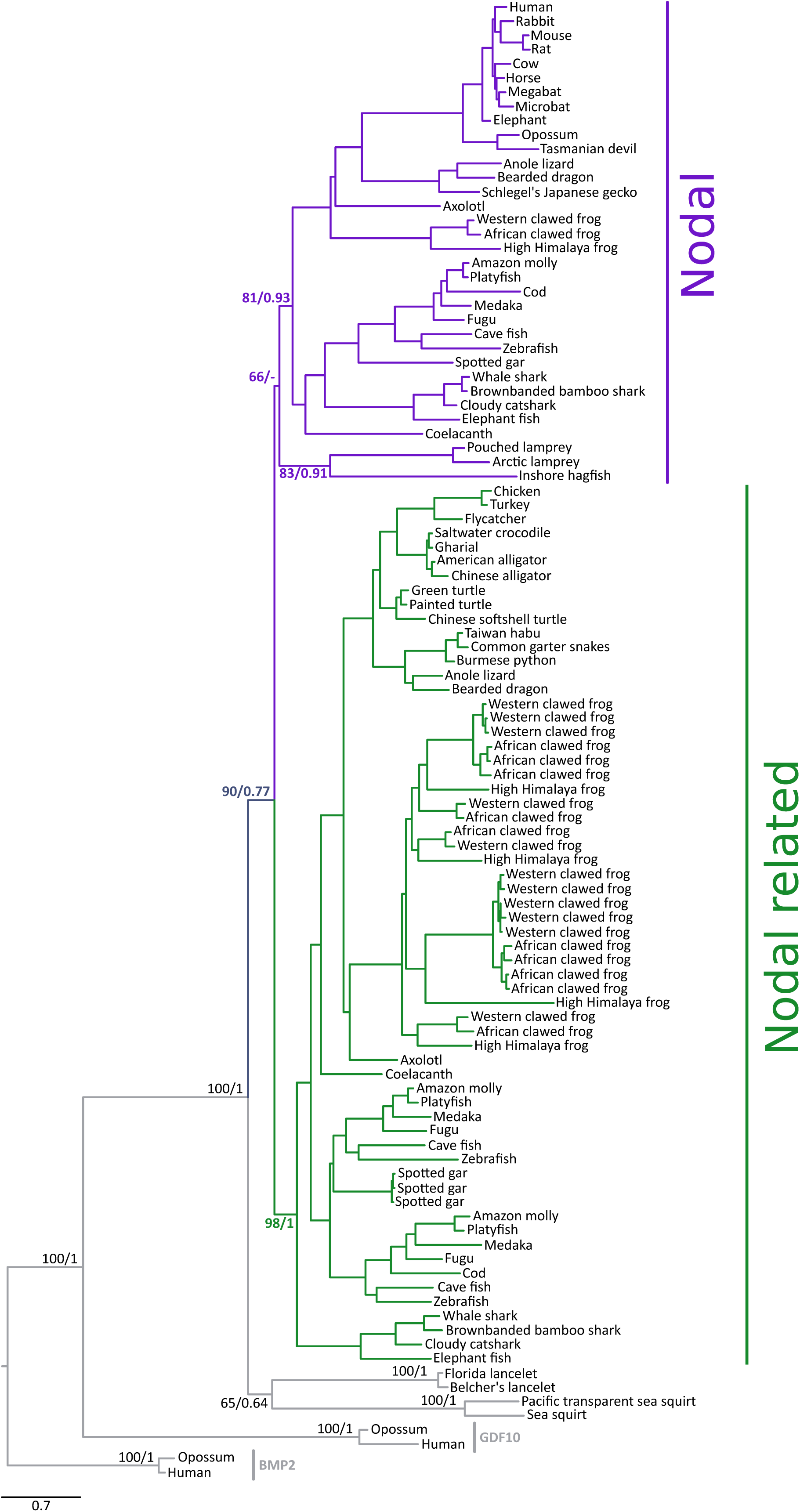
A maximum-likelihood tree depicting evolutionary relationships among the Nodal and Nodal-related genes of chordates. Numbers on the nodes represent maximum likelihood ultrafast bootstrap and Bayesian posterior probability support values. BMP2 sequences from human (*Homo sapiens*) and opossum (*Monodelphis domestica*) were used as outgroups.

The results of our synteny analyses are consistent with the phylogenetic results: the Nodal and Nodal-related genes are each in a relatively well-conserved chromosomal location (Fig. 2). In almost all vertebrates, Nodal is flanked by an ortholog of EIF4EBP2 upstream and of PALD1 downstream, and Nodal-related is flanked by an EIF4EBP1 ortholog upstream and orthologs of ASH2L and STAR downstream (Fig. 2). In both cases, there is higher synteny conservation among bony vertebrates, from human to spotted gar. Interestingly, there is high synteny conservation for the regions corresponding to the Nodal and Nodal-related genes even in the lineages where these Nodal genes have been lost. Synteny comparisons between cyclostomes and the Nodal or Nodal-related genes of gnathostomes were uninformative.

**Figure 2.**
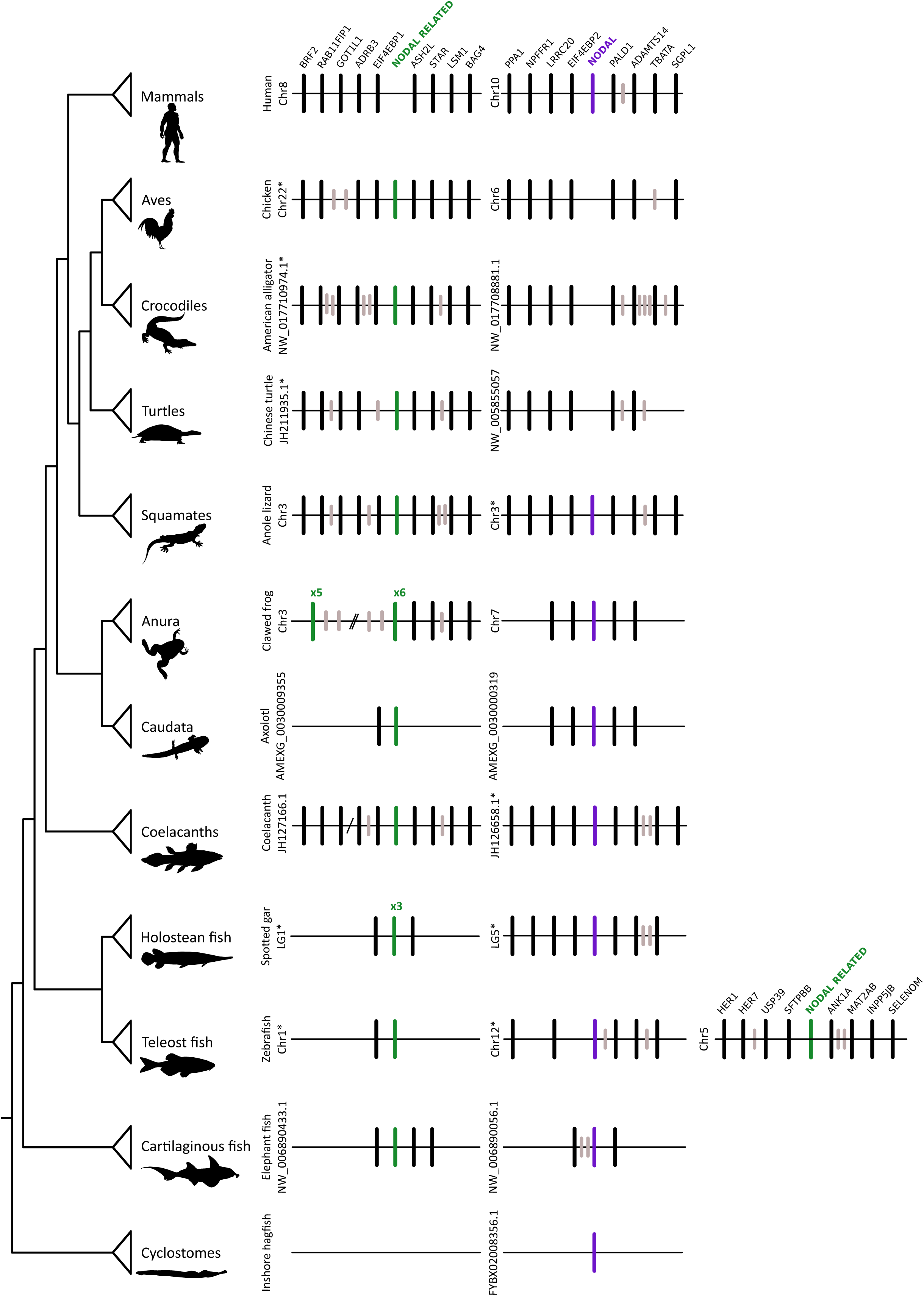
Gene order in the chromosomal region containing the Nodal and Nodal-related genes of vertebrates. Asterisks denote that the orientation of the genomic piece is from 3’ to 5’, gray lines represent intervening genes that do not contribute to conserved synteny.

### 3.2 Evolution of Nodal and Nodal-related repertoires in fish and amphibians

The cases of fish and amphibians deserve more careful attention, as these groups present additional variation in copy number of Nodal and Nodal-related genes relative to other vertebrates. In both lineages, Nodal has remained as a single-copy ortholog, despite inconsistencies in the nomenclature. For example, the Nodal genes of fish fall in a clade (Figs. 1 and 3) and include the single ortholog gene cyclops, also known as ndr2, for nodal-related-2, of zebrafish. The three copies of the Nodal-related gene from spotted gar are in an exclusive clade (Fig. 1 and 4), and the teleost Nodal-related genes are grouped into two clades, one of which includes the southpaw gene of zebrafish, also known as ndr3, and is sister to the clade that includes the three spotted gar paralogs and the other that includes the squint gene of zebrafish, also known as ndr1 (Figs. 1 and 4). Thus, most teleosts have a single copy of Nodal, and duplicate copies of Nodal-related, whereas the spotted gar has a single copy of Nodal, and triplicate copies of Nodal-related.

**Figure 3.**
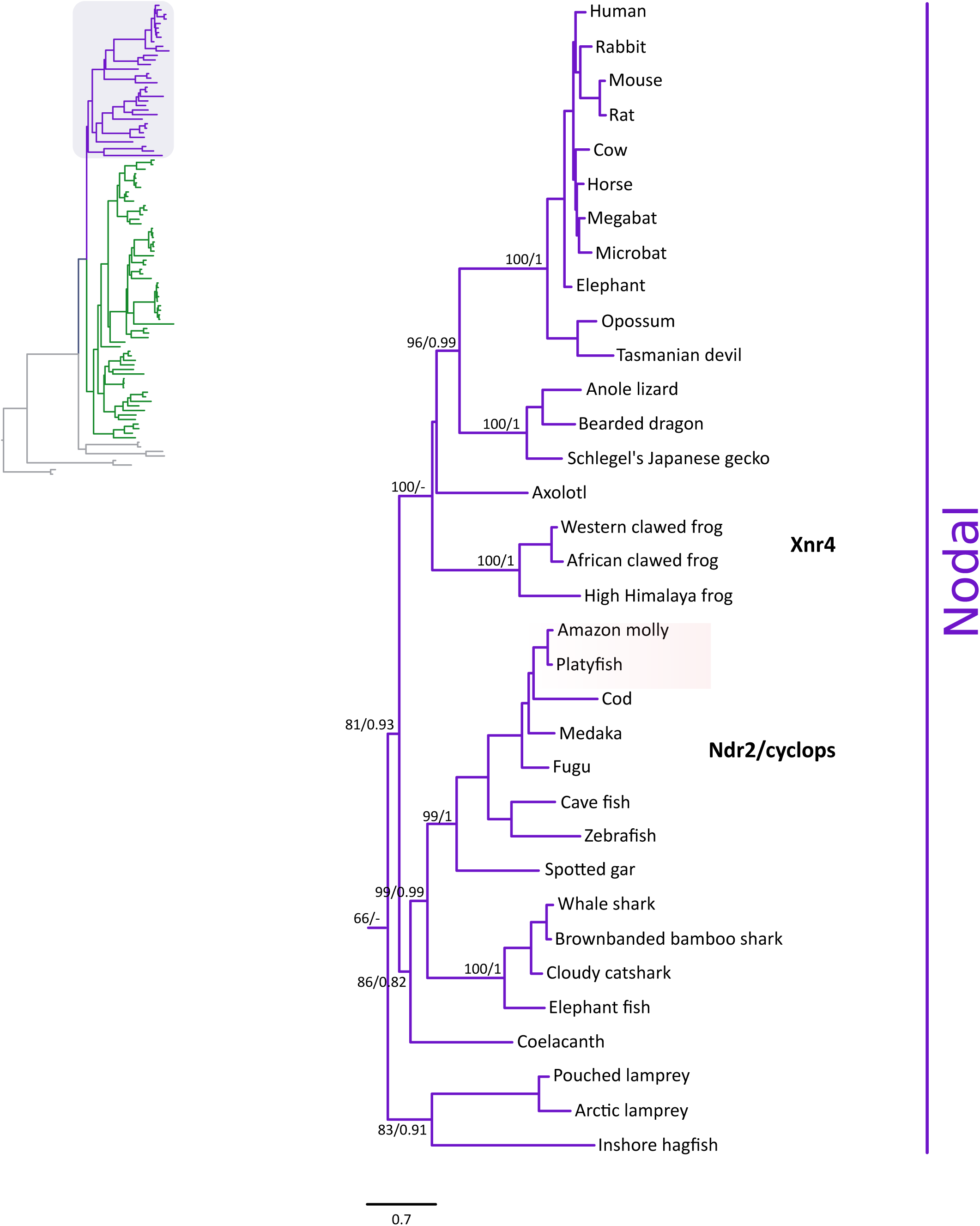
Maximum likelihood tree depicting evolutionary relationships among Nodal genes in vertebrates. Numbers on the nodes correspond to maximum likelihood ultrafast bootstrap support and Bayesian posterior probability support values. This tree topology does not represent a novel phylogenetic analysis; it is the Nodal clade that was recovered from Fig. 1.

**Figure 4.**
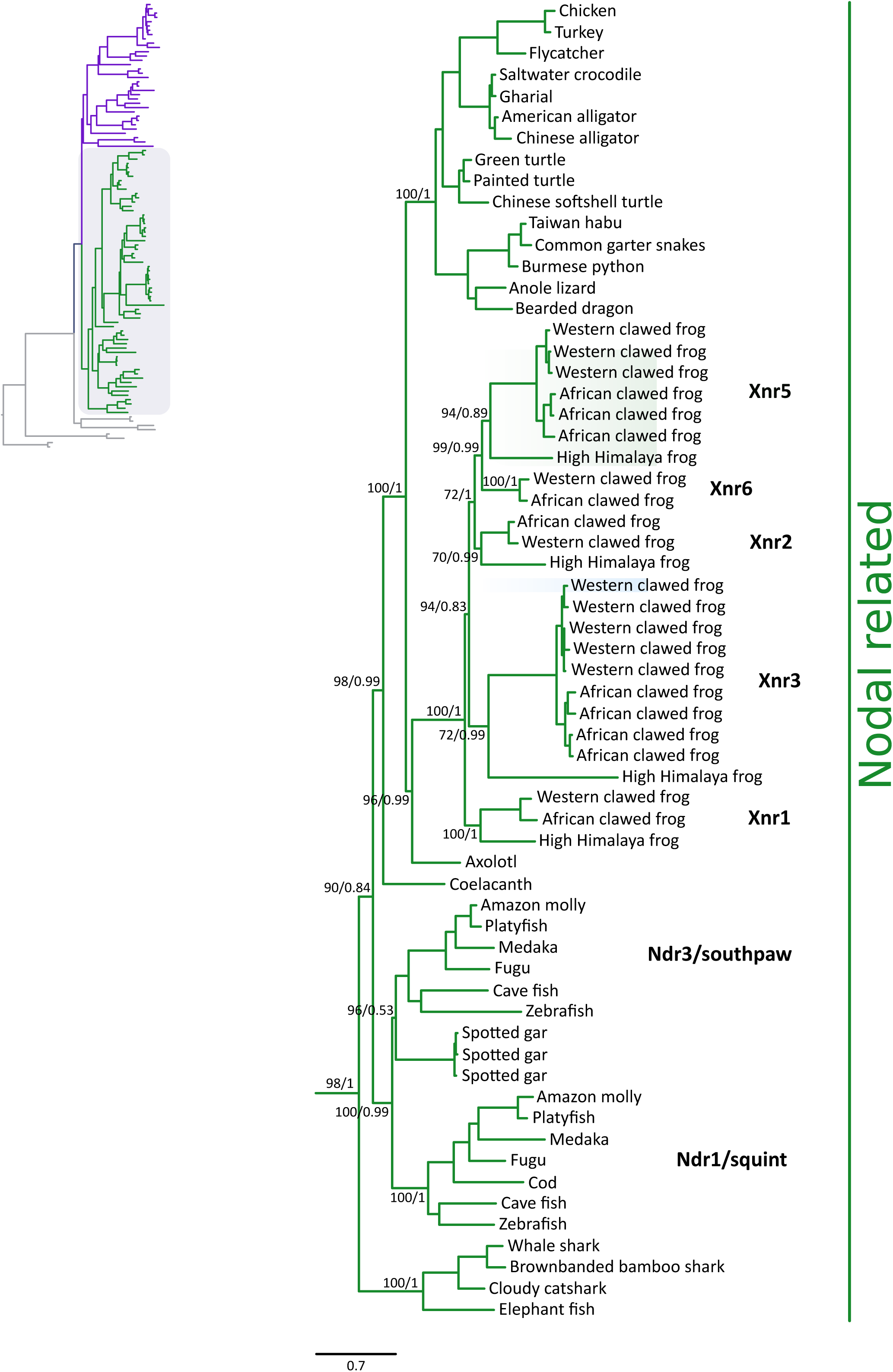
Maximum likelihood tree depicting evolutionary relationships among Nodal-related genes in vertebrates. Numbers on the nodes correspond to maximum likelihood ultrafast bootstrap support and Bayesian posterior probability support values. This tree topology does not represent a novel phylogenetic analysis; it is the Nodal clade that was recovered from Fig. 1.

The amphibian repertoire is the most variable among the groups studied. Axolotl, order Caudata, and frogs, order Anura, include both Nodal and Nodal-related genes in their genomes. There is a single ortholog of Nodal in all amphibians, known as Xnr4, for Xenopus-nodal-related-4. By contrast, there is a single copy of Nodal-related in axolotl, but three to eleven copies in anurans, including the Xnr1, 2, 3, 5 and 6 genes. The three frogs in our study include a single ortholog of Xnr1, which is sister to all other Nodal-related genes from frogs (Fig. 4), and a single ortholog of Xnr2. Xnr6 is also present as a single copy in the two clawed frogs but is absent from the current assembly of the High Himalaya frog. There are single copies of Xnr3 and 5 in the High Himalaya frog, but these two genes have expanded independently to 3 to 5 copies in the Western and African clawed frogs respectively (Figs. 1 and 4).

## 4 DISCUSSION

We studied the evolution of Nodal and Nodal-related genes with the main goal of integrating their evolutionary history with that of the GDF1/3 gene family to gain insights into the evolution of the heterodimer that is responsible for triggering the Nodal pathway early in development. Further, understanding the evolutionary history and phyletic distribution of these genes is also important as Nodal is expressed in multiple types of cancers (Strizzi et al., 2012) and the nodal pathway has emerged as a potential target for cancer therapy (Kalyan et al., 2017). Thus, our inferences of orthology should facilitate the selection of adequate model organisms for this research, and permit an evolutionarily sensible interpretation of experimental outcomes.

### 4.1 Evolution of the Nodal and Nodal-related genes in jawed vertebrates

The phylogenetic and synteny analyses resolve orthology relationships and the duplicative history of the Nodal and Nodal-related genes of vertebrates, placing them in reciprocally monophyletic groups (Fig. 1). Previous studies had identified a discordant number of gene lineages, probably because of scarce sequence information outside mammals and actinopterygian fish (Fan and Dougan, 2007; Kuraku and Kuratani, 2011). The key difference among these studies lied in the position of the clade that includes the teleost ndr2/cyclops genes. With the more extensive taxonomic sampling of our study, we are able to resolve the evolutionary history of the Nodal and Nodal-related genes of jawed vertebrates to two well-supported gene lineages, Nodal and Nodal-related (Fig. 1).

### 4.2 Inference of ancestral repertoires

Because both Nodal and Nodal-related are present in cartilaginous fish, we infer that these genes were present at least in the ancestor of jawed vertebrates, between 615 and 473 mya. Synteny is relatively well conserved across jawed vertebrates (Fig. 2), and the synteny blocks of Nodal, 8.1, and Nodal-related, 10.3, in human map to chromosomal segments that derive from the same ancestral chordate linkage group (CGL6 from Putnam et al. 2008). Thus, we infer that Nodal and Nodal-related derive from the whole genome duplications early in vertebrate evolution. In agreement with this, most non-vertebrate species screened maintain a single ortholog of the gene (Grande et al., 2014). Interestingly, we were able to find Nodal-like genes in the current genome assemblies of both hagfish and lampreys, the two extant lineages of cyclostomes, whereas a previous study included no Nodal-like gene in a particular lamprey species, the sea lamprey (*Petromyzon marinus*) (Lagadec et al., 2015). In our results, phylogenetic analyses would suggest the orthology of the identified cyclostome Nodal-like gene to the Nodal gene of jawed vertebrates, rather than Nodal-related, although the support is not high (Figs. 1 and 3).

Taken together, our results indicate that the duplication of Nodal and Nodal-related genes can be traced back at least to the last common ancestor of jawed vertebrates, and possibly to the last common ancestor of extant vertebrates. Unfortunately, as in other gene families such as globins, Myb and KCNA, resolving orthology between cyclostome and gnathostomes remains a hard problem because of GC-, codon-, and amino acid composition biases (Qiu et al., 2011; Smith et al., 2013; Schwarze et al., 2014; Campanini et al., 2015; Opazo et al., 2015). This has complicated inferences of the phenotype of the last common ancestor of vertebrates on the basis of gene orthology (reviewed in (Onimaru and Kuraku, 2018)).

### 4.3 Differential retention of Nodal and Nodal-related genes

Tracing gains and losses along the vertebrate phylogeny shows that the differential retention of ancestral genes has played an important role in shaping Nodal repertoires (Fig 5). Nodal is present in all major lineages of vertebrates other than birds, crocodiles, turtles, and snakes (Fig. 3), suggesting that it was lost twice independently, that is, in the common ancestor of archelosaurians (birds, crocodiles, and turtles), between 280 and 254 millions of years ago, and in the snake lineage. On the other hand, Nodal-related is present in all main groups of vertebrates other than mammals and cyclostomes (Fig. 4).

**Fig. 5.**
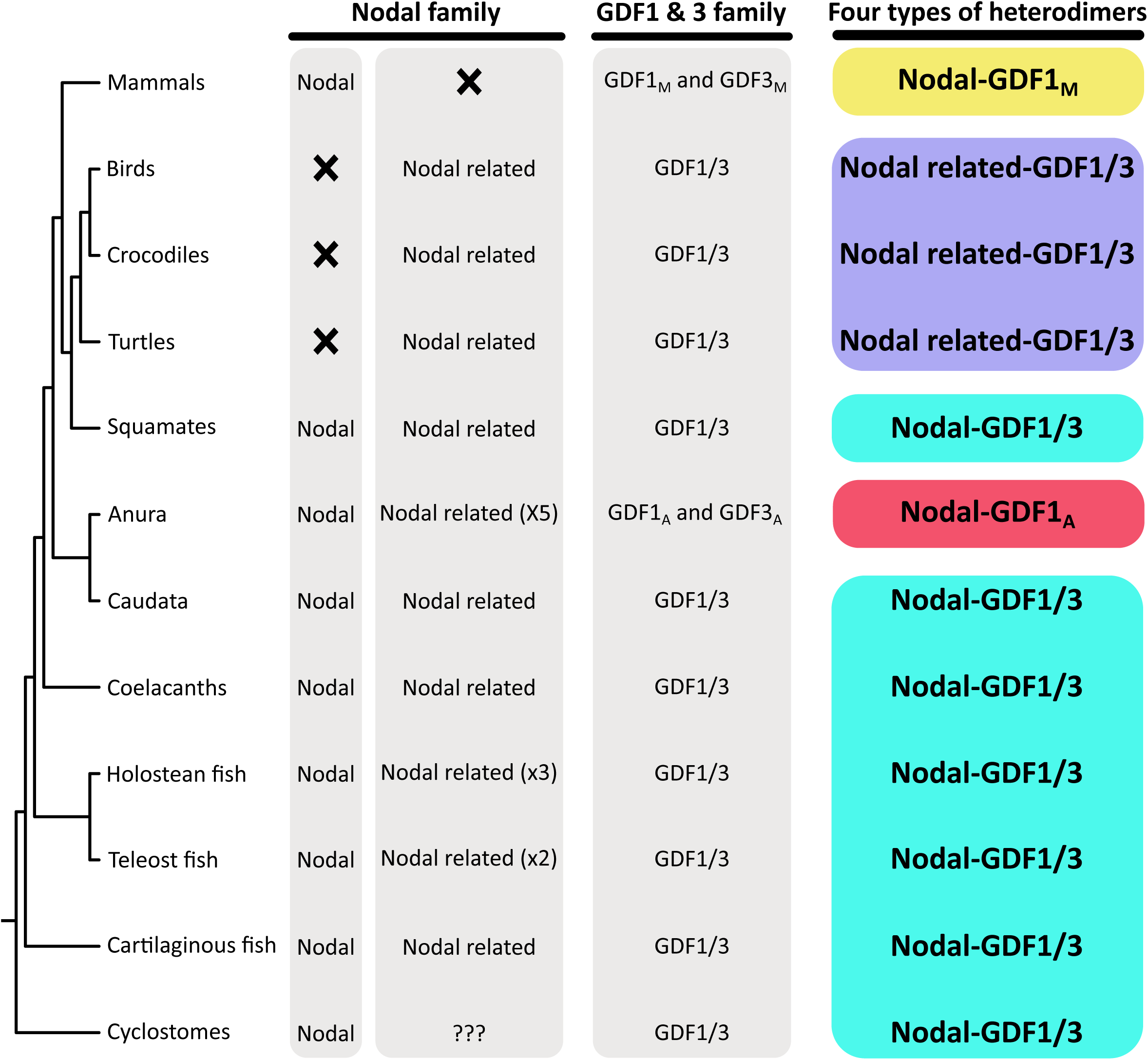
Phyletic distribution of Nodal, Nodal-related, GDF1/3, GDF1 and GDF3 genes among vertebrates, including the putative composition of the heterodimer. The different colors refer to the putative four types of heterodimers that can trigger the Nodal pathway in vertebrates. The phyletic distribution of the GDF1/3, GDF1 and GDF3 genes was obtained from Opazo & Zavala (2018).

The case of teleost fish deserves attention given that this group experienced an additional whole genome duplication in their early evolution (Amores et al., 1998; Petit et al., 2004; Kuraku and Meyer, 2009). Accordingly, we expect a more diverse repertoire in comparison to other vertebrates. This was the case, as teleosts have two Nodal-related genes: ndr1/squint and ndr3/southpaw, and one Nodal ortholog, ndr2/cyclops. Fan & Dougan (2007) suggested that the diversity of Nodal genes in teleosts arouse as a combination of the teleost-specific genome duplication and small-scale duplication events. According to these authors, the teleost-specific genome duplication would have given rise to two ancestral genes. Later, one of them would have undergone a duplication event giving rise to the ndr1/squint and ndr3/southpaw genes (Fan and Dougan, 2007). By contrast, our results indicate that the ancestor of teleosts already had two genes, which as a product of the teleost-specific genome duplication would have given rise to a repertoire of four. After this genome duplication, both Nodal-related ohnologs were retained (ndr1/squint and ndr3/southpaw)(Figs. 1 and 4), whereas only one of the Nodal ohnologs was retained (ndr2/cyclops)(Figs. 1 and 3).

We observed additional lineage-specific expansions of the Nodal-related gene in the spotted gar and amphibians of the order Anura, the group that includes frogs and toads (Fig. 4). The spotted gar has three Nodal-related genes (Figs. 1 and 4) which are very similar among them with pairwise distances ranging from 2.43% to 3.78%, and group with the ndr3/squint gene of teleosts. The most important gene expansion occurred in anurans. The duplications giving rise to the Xnr1, 2, 3, 5 and 6 genes are shared among all frogs examined. The Xnr6 has been apparently lost in the High Himalaya frog, and the Xnr3 and Xnr5 genes have expanded further in the two clawed frog lineages independently. Thus, our analyses indicate that the expansion of Nodal-related genes occurred in two waves, the first was after the divergence of the orders Anura and Caudata but before the radiation of anuran species, whereas the second wave happened independently in the western and African clawed frog species (Fig. 4).

### 4.4 Gene duplication and gene loss as sources of evolutionary innovations

The differential retention of genes could be seen as a stochastic process, in which the resulting differences in gene complement do not translate into functional consequences. This could be due to some functional redundancy among the paralogs that could work as a backup (i.e., functionally overlapping paralogues) in the case one is lost or inactivated (Cañestro et al., 2009; Félix and Barkoulas, 2015; Albalat and Cañestro, 2016), or a by-product of divergent nature of the genomic location in which the duplicates are found (Hara et al., 2018b, 2018a). Alternatively, it is also possible that the presence of multiples copies opens opportunities for the emergence of biological novelties (Ohno et al., 1968; Ohno, 1970; Hughes et al., 1994; Force et al., 1999; Zhang, 2003). For example, the differential retention of functional copies of the INSL4 gene in catarrhine primates has been associated to unique reproductive traits of the group (Arroyo et al., 2012a, 2012b; Malone et al., 2017).

The loss of Nodal in birds is informative in this regard because several studies have reported the expression of the Nodal gene during early development of the chicken (Raya et al. 2004a,b, and references therein). Thus, we infer these studies were actually reporting the expression of the Nodal-related gene, which is annotated as Nodal (https://www.ncbi.nlm.nih.gov/gene/395205, NCBI). This is an example of how ‘hidden paralogy’, the differential retention of alternate paralogs, has complicated reconstructions of ancestral states in vertebrate evolution (reviewed in <u>Kuraku 2010</u>) and highlights the need for applying phylogenetically explicit approaches to make orthology inferences (Gabaldón, 2008; Altenhoff et al., 2016). From a functional standpoint, the Nodal-related gene of birds seems to be fulfilling the role of Nodal (Raya et al. 2004a,b, and references therein), suggesting these paralogs have retained some degree of redundancy.

### 4.5 Evolution of the Nodal-GDF1/3 heterodimers

There has been evidence for the co-dependence of the Nodal and GDF1/3 gene families during early developmental processes in all main group of vertebrates for some time, but their interactions were not clear. This has recently changed, as three independent groups using different approaches reported that the formation heterodimers between Nodal and GDF1/3 is necessary to activate the Nodal pathway (Bisgrove et al., 2017; Montague and Schier, 2017; PellicciaJ.L. et al., 2017).

From an evolutionary standpoint, the vertebrate Nodal/Nodal-related and GDF1/3 gene families have undergone lineage-specific expansions and retentions in different vertebrate lineages (Opazo and Zavala, 2018). As a result, the signaling heterodimer, which triggers the Nodal developmental pathway, is composed by proteins that derive from different genes in different vertebrate groups (Fig. 5). Mapping the presence of the corresponding genes indicates that cyclostomes, cartilaginous fish, teleosts, holostean fish, caudates, and lizards possess a single ortholog of Nodal and a single ortholog of GDF1/3 gene (Fig. 5 and 6). So, these groups would be able to form heterodimers following the conformation recently reported (Bisgrove et al., 2017; Montague and Schier, 2017; PellicciaJ.L. et al., 2017). In birds, turtles, crocodiles, and snakes, however, there is no Nodal gene, as far as the species analyzed so far are concerned. Evidence from chicken suggests that Nodal-related is involved in forming the heterodimers with GDF1/3 (Raya et al., 2004a, 2004b), and this is probably the case for all the groups listed above, in a case that illustrates the evolutionary significance of functional redundancy.

The case of anurans and mammals is also interesting; as they both possess a single ortholog of Nodal and independently derived paralogs of GDF1/3, the GDF1_A_ and GDF3_A_ genes in amphibians derive from an amphibian-specific duplication whereas the GDF1_M_ and GDF3_M_ genes of mammals derive from a mammalian-specific duplication (Fig. 5) (Opazo and Zavala, 2018). Thus, the Nodal genes of mammals and anurans are 1:1 orthologs, but their partners for the heterodimer, GDF1 or GDF3, are not (Opazo and Zavala, 2018). Thus, mammals activate the pathway with a heterodimer containing Nodal and GDF1_M_ (Fig. 5), whereas anurans trigger the Nodal developmental pathway with Nodal and GDF1_A_ (Vg1)(Fig. 5).

In summary, there are at least four different heterodimer conformations that activate the Nodal pathway in vertebrates (Fig. 5): 1) Nodal + GDF1/3, in cyclostomes, cartilaginous fish, teleosts, holostean fish, caudates and lizards; 2) Nodal-related + GDF1/3, in birds, crocodiles, turtles, and snakes; 3) Nodal and GDF1_A_ in anurans; and 4) Nodal + GDF1_M_ in mammals. The diversity of heterodimers that can trigger the same developmental pathway highlights the redundancy present in the genomes of vertebrates and the versatility of the evolutionary process. In this case, the duplication that gave rise to Nodal and Nodal-related predates the divergence between cartilaginous fish and the rest of extant jawed vertebrates (615-473 mya), and one of the duplicates, Nodal, was lost in the common ancestor of archelosaurs (312-254 mya). Despite their relatively old divergence, both nodal and nodal-related seem to have retained the ability to activate the pathway in the absence of nodal, suggesting that functional redundancy can last long periods of time even if not directly needed. Furthermore, it is important to note that the role of Nodal-related in species that also have Nodal is poorly understood, which suggests that these species could form a much more diverse array of heterodimers. Thus, comparing the Nodal pathway in representative species belonging to the four groups of heterodimer conformations will shed light regarding the robustness of the nodal pathway.

## ACKNOWLEDGMENTS

This work was supported by the Fondo Nacional de Desarrollo Científico y Tecnológico (FONDECYT 1160627) grant to JCO. FGH’s work was partially funded by the National Science Foundation through grant EPSCoR RII Track-2 FEC 1736026.

## SUPPORTING INFORMATION

Additional supporting information may be found online in the Supporting Information section at the end of the article.

